# Exotic and Zoological Birds Resident and Imported into Nigeria harbour Highly Pathogenic Avian Influenza Virus: Threat to Poultry Production, Food security and Public Health

**DOI:** 10.1101/2025.01.22.634354

**Authors:** Atuman Yakubu, Bitrus Inuwa, Judith Bakam, Edmond Orioko, Hambolu Sunday Emmanuel, Clement Meseko

**Affiliations:** National Veterinary Research Institute, Vom, Nigeria; Department of Veterinary Services, Ministry of Agriculture, Delta State, Nigeria

**Keywords:** HPAI, Exotic and wild birds, Importation, LBMs, Parks and Gardens, Nigeria

## Abstract

Influenza is a major infectious disease challenge affecting animal and human health globally, and wild birds are historically the primary reservoirs of all the known Influenza A virus subtypes. Here, we detected the Highly Pathogenic Avian Influenza (HPAI) virus in exotic and aquatic birds in three different locations in Nigeria. On the 8^th^ of February 2021, exotic birds: Yellow Golden Pheasant (*Chrysolophus pictus*), Sultan chicken (*Gallus gallus domesticus)*, Lakenvelder chicken (*Gallus gallus domesticus*), and Common pheasant (*Phasianus calchicus*), imported from Libya and transported across the Niger Republic border to Nigeria, were presented to the National Veterinary Research Institute, Vom, for screening. Also, a family in Lagos State bought some exotic aquatic birds from a live bird market in Sokoto State, Nigeria, where sudden death was recorded with the birds showing few clinical signs. Similarly, the sudden death of some aquatic birds was reported in Mandela Parks and Gardens in Asaba, Delta State, few weeks after some captured wild birds were introduced to the Park and Gardens. Oropharyngeal, cloacal, and tissue samples were all collected from the reported cases. Total viral nucleic acid was extracted and screened for Influenza A viruses using real-time RT-PCR. The HPAI viruses H5N1 and H5N8 were detected in the imported aquatic (geese and ducks) and exotic (yellow golden pheasant) birds. The samples tested negative for low-pathogenic Avian Influenza Virus (H9N2) as well as other avian viruses, viz., *Avian avulavirus-1 (*Newcastle disease Virus) and infectious bronchitis virus. This highlights the role of these resident and imported exotic birds in the local transmission and spread of the HPAI virus to domestic poultry. The findings call for proper biosecurity and quarantine measures for exotic and wild birds to reduce the potential risk to animal and public health in Nigeria.

## 1. Introduction

Influenza A is among the major infectious disease problems affecting animal and human health globally (Kuchipudi and Nissly, 2018). In 1996, a strain of Highly Pathogenic Avian Influenza (HPAI) H5N1, termed the Goose/Guangdong (Gs/GD) virus, emerged in Chinese poultry and has since evolved into different clades and multiple sub-clades that continue to threaten poultry production and food security, especially in developing countries (Abolnik *et al*., 2022). Influenza A virus (IAV) is classified within the *Alphainfluenza* genus of the *Orthomyxoviridae* family as an enveloped single-stranded, negative-sense RNA virus that comprises eight genome segments, two of which code for the glycoproteins heamagglutinin (HA) and neuraminidase (NA), whose antigenic properties are used to classify viral subtypes (McCauley *et al*., 2019). To date, sixteen haemagglutinin (HA) and nine neuraminidase (NA) virus subtypes have been described in avian species, and almost all known subtype combinations have been isolated from wild birds (Tong *et al*., 2012). Avian influenza outbreak is always a concern due to its economic impact and public health threat. In Nigeria, the first outbreak of HPAI was reported in 2006, which caused huge economic losses to poultry farmers. The introduction of HPAI in Nigeria was attributed to the activities of migratory birds and trade in poultry products as possible sources of infection (Meseko *et al*., 2010; Twinning, 2021). The first avian influenza virus (AIV) outbreak was successfully contained; however, the recurrent reports of HPAI outbreaks in poultry farms and live bird markets (LBMs) in some parts of the country underscore the speculations about the possible role of wild birds in the maintenance and spread of infection (Coker *et al*., 2014). In January 2015, Nigeria again confirmed the presence of H5N1 HPAI in poultry. This was the first occurrence of H5N1 HPAI in the country since the last epidemic between 2006/2008 and 2015/2016 (FAO, 2015; Monne *et al*., 2015). Till date, since the first outbreak of AIV in 2006 in Nigeria, five subtypes of HPAI and LPAI (H5N1, H5N2, H5N6, H5N8, and H9N2) have been reported (Meseko *et al*., 2010; Shittu *et al*., 2020; Sulaiman *et al*., 2021; Ameji *et al*., 2022; Laleye *et al*., 2022; Meseko *et al*., 2023). In 2021, a new outbreak of AIV was reported in upto thirty states across Nigeria, with a novel strain of HPAI H5N1/H5N8 clade 2.3.4.4b being introduced into the country. And as at December 2022, over 400 positive cases of the HPAI was recorded (Meseko *et al*, 2023). The re-introduction and spread of HPAI were linked to wild birds and waterfowls in backyards and live bird markets. The Wild birds of the orders Anseriformes (ducks, geese, and swans) and Charadriiformes (gulls and shorebirds) are the historically natural reservoirs of influenza A viruses and have since played an important role in the maintenance and global spread of AIV. Continued monitoring and screening of AIV activity in wild exotic and aquatic birds not normally symptomatic (Meseko *et al*., 2010; Umar *et al*., 2023; Olawuyi *et al*., 2023) may improve risk assessment for poultry producers and early detection to curtail further spread. In this study, we screened for AIV healthy exotic birds imported across the border to Nigeria and also investigated the cause of mortality in some aquatic birds in recreational gardens and parks after some captured wild birds were introduced into the bird holding.

## 2.0 MATERIALS AND METHODS

### 2.1 Case History

On the 8^th^ of February 2021, exotic birds: the Yellow Golden Pheasant (*Chrysolophus pictus*) *n =* 2, Sultan chicken (*Gallus gallus domesticus*) *n =* 2, Lakenvelder chicken (*Gallus gallus domesticus*) *n =* 1, and Common Pheasant (*Phasianus colchicus*) *n =* 1 (Figure 1 a-e) were imported from Libya across the Niger Republic border to Nigeria. These birds were presented to the National Veterinary Research Institute NVRI, Bauchi State, for routine screening. On the 30^th^ of April 2021, a family in Lagos State, Nigeria, bought 2 peacocks, 4 geese, and some imported ducks (*n =* 3) from a live bird market (LBM) in Sokoto State, where two of the geese and one of the ducks died without showing any clinical signs few days after their arrival. The peacock died after showing clinical signs of wobbling feet and uncoordinated movement. Similarly, the mortality of 3 ducks, 1 peacock and, 1 ostrich was reported from Mandela Parks and Gardens in Asaba, Delta State, in August 2021, a few weeks after some captured wild birds were introduced to the park and gardens (Figure f-h).

**Figure 1:**
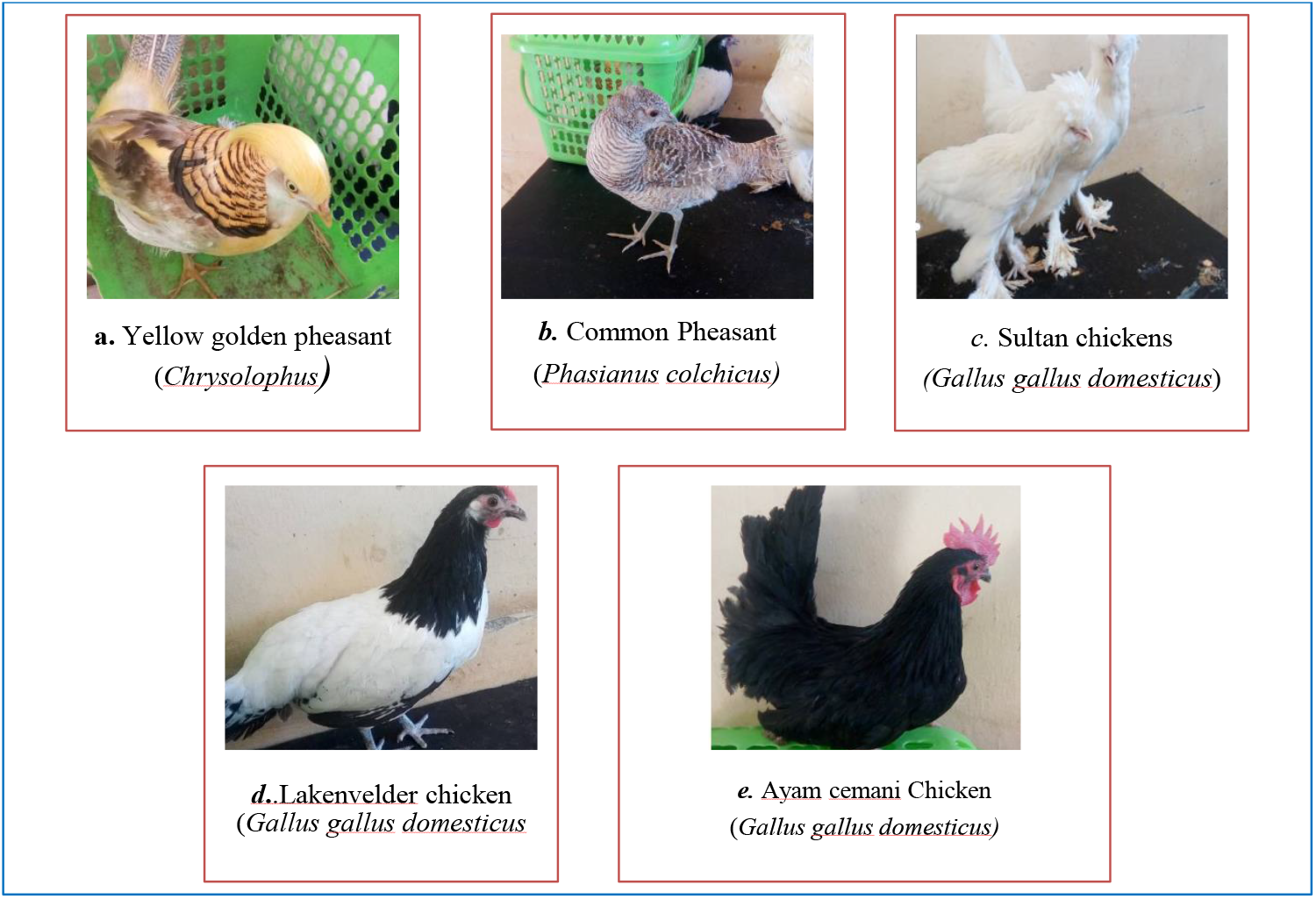
(a-e). Birds Imported to Nigeria from Niger Republic.

**Figure 2:**
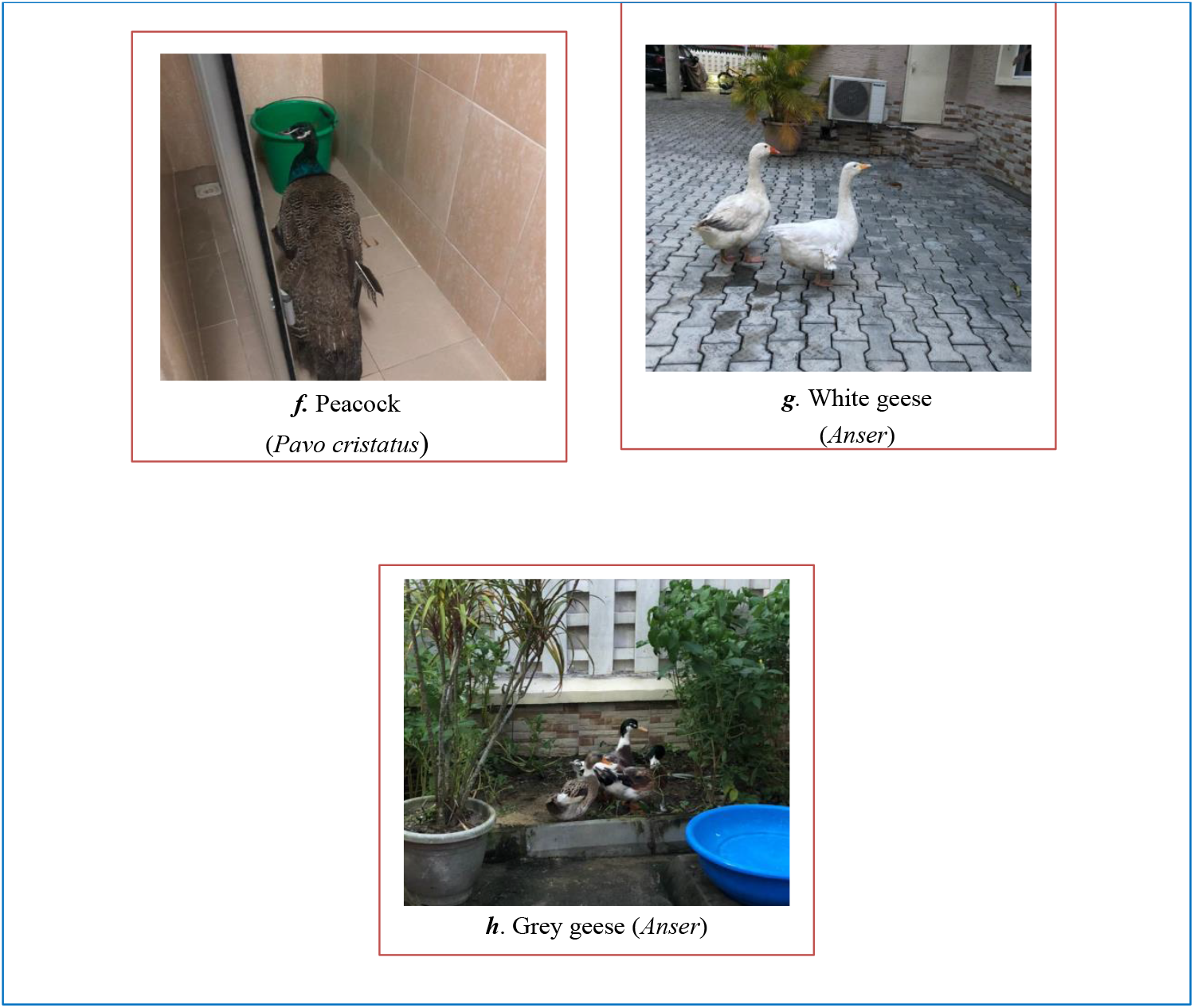
(f-h). Some imported Birds from Sokoto LBM and Mandela Parks and Gardens, Asaba, Delta State.

### 2.2 Samples Collection and Processing

Oropharyngeal and cloacal swab samples were collected in a 1 ml viral transport medium (VTM) from the exotic birds presented for routine screening on importation from Libya. Tissue samples were also collected at postmortem from death carcasses from Lagos and Asaba, Delta States, respectively. All samples were properly labeled and transported in a cooler on ice packs to the Regional Laboratory for Animal Influenza and Transboundary Disease Laboratory, NVRI Vom, Nigeria. All samples were processed and kept at -80°C freezer until testing.

### 2.3 Nucleic Acid Extraction

Total viral nucleic acid was extracted using the Da An Gene RNA/DNA purification kit (Da An Co., Ltd.), following the manufacturer’s instructions. Nucleic acids were eluted in 50 µl elution buffer and stored at -20ºC until further analysis.

### 2.4 RT-PCR

Extracted viral nucleic acids were first screened for the Avian Influenza Virus (AIV) Matrix gene using the Qiagen® QuantiTect Multiplex RT-PCR Kit, as previously described by Spackman *et al*. (2002). This protocol amplifies different regions within the matrix (M) gene, which is well conserved among avian influenza viruses of all subtypes. All samples that tested positive for the AIV matrix gene were subsequently screened for HPAI H5N1, H5N8 and LPAI H9N2 as previously described (Slomka *et al*., 2007; Monne et al., 2008; Hoffman *et al*., 2016), as well as other common endemic avian viruses, including *Avian avulavirus*-1 (AOV-1) and infectious bronchitis virus (IBV) in the region using protocols previously reported that detect the viral genome for AOV-1 and IBV respectively (Callison *et al*., 2006; Fuller *et al*., 2009; Shittu *et al*., 2018; Bitrus *et al*., 2020).

## 3. Results

The avian Influenza matrix gene was detected in one out of five apparently healthy exotic birds (Yellow Golden Pheasants) imported from Libya and presented for screening (Table 1). After further screening with primers and probes specific for the haemagglutinin and neuraminidase genes, the Highly Pathogenic Avian Influenza Virus of the subtype H5N8 was detected (Table 1). While the other two species, Sultan birds and Japanese Phantom, were all negative for the AIV matrix gene (Table 1). The AIV matrix gene was also detected in two of the geese and one of the ducks purchased by a family in LBM in Sokoto and moved to Lagos. HPAI of the subtype H5N1 was further confirmed in these samples. The HPAI H5N1 was also detected in one of the duck samples submitted from the Mandela Park and Garden, Asaba, Delta State. All samples tested negative for LPAI (H9N2), Newcastle disease virus (NDV), and Infectious bronchitis (IBV) (Table 1).

**Table 1:** Highly Pathogenic Avian Influenza Virus Detected in Exotic and Aquatic Birds in Nigeria.

| Source of birds | Exotic birds | Samples collected | Mortality | AIV subtype detected |  | Other common avian viral diseases tested |  |
| --- | --- | --- | --- | --- | --- | --- | --- |
|  |  |  |  | HPAI | LPAI |  |  |
|  |  |  |  |  |  | NDV | IBV |
| From Libya through Niger Republic border to Bauchi State, Nigeria | Yellow Golden Pheasant | 2 | 0 | H5N8 | 0 | 0 | 0 |
|  | Sultan chicken | 2 | 0 | 0 | 0 | 0 | 0 |
|  | Common Pheasant | 1 | 0 | 0 | 0 | 0 | 0 |
|  | Lakenvelder chicken | 1 | 0 | 0 | 0 | 0 | 0 |
| From LBM Sokoto to Lagos State | Peacock | 2 | 0 | 0 | 0 | 0 | 0 |
|  | Geese | 4 | 2 | H5N1 | 0 | 0 | 0 |
|  | Duck | 3 | 1 | H5N1 | 0 | 0 | 0 |
| Captured wild birds introduced into Mandela Park and Gardens, Asaba, Delta State | Peacock | 1 | 1 | 0 | 0 | 0 | 0 |
|  | Ostrich | 1 | 1 | 0 | 0 | 0 | 0 |
|  | Ducks | 3 | 2 | H5N1 | 0 | 0 | 0 |

## 4. Discussion

The highly pathogenic avian influenza (HPAI) virus has continued to ravage the poultry industry globally, causing huge socio-economic losses associated with the high mortality in poultry, culling and depopulation of birds for control, and loss of livelihood for farmers and farm workers in Nigeria, where the poultry industry plays a vital role in food security. The detection of HPAI H5N1 and H5N8 in the three different locations in Nigeria in this study has confirmed previous studies on the role of captive-wild birds and ducks in the epidemiology and the continued spread of the virus in Nigeria (Joannis *et al*., 2008; Ameji *et al*., 2022; Olawuyi *et al*., 2023), as reported in other parts of the world such as Cambodia (Desvaux *et al*., 2009); Bangladesh (Hassan *et al*., 2017); Iran (Fallah *et al*., 2020); and India (Nagarajan *et al*., 2017). In 2021, an outbreak of AIV was reported in more than 20 states across the six geographical zones of the country with the new introduction of a novel strain of HPAI H5N1/H5N8 clade 2.3.4.4b (Meseko *et al*., 2023).Clade 2.3.4.4b has been circulating since and has become one of the most serious threats for both poultry industries and smallholder farmers worldwide (Abolnik *et al*., 2019). The HPAI H5N8 was detected in an apparently healthy exotic bird (Yellow Golden Pheasant) imported to the country through the Republic of Niger from Libya, the exotic birds may not be rule out in the spread, as recent data from Libya (Kammon *et al*.,. 2022) and Niger Republic (FAO Report, 2021) further indicate AIV occurrences. The trade of captive wild birds such as Yellow Golden Pheasant, Sultan bird, Lakenvelder, and Japanese Phantom is on the increase in Nigeria. Illegal trade, poor border and inter-State surveillance of these exotic birds may continue to play a greater role in the ecology and epidemiology of AIVs in the country. However, given that these exotic birds have been understudied in comparison to other types of species of birds, their potential importance in the natural history of AIVs in the country remain unknown. The detection of HPAI H5N1 in a dead duck and geese purchased from LBMs in Sokoto and transported to Lagos State in this study has confirmed the report of Sulaiman *et al*., (2021), that LBM as a potential source of the IAV viruses. Naive birds in LBMs frequently get into contact with infected birds. The newly infected birds can be sold or returned to the same farm, allowing AIV to spread within LBMs as well as from LBMs to farms in different geographical areas (Pepin *et al*, 2013). The HPAI H5N1 virus resulted in the death of birds in a park and gardens when captured wild birds were introduced into the garden observed in this study. Parks and gardens in Nigeria are majorly recreational, educational, and tourist centers where human activities take place on a daily basis. Aside from the socioeconomic impact, IAV has zoonotic potential as all human influenza pandemics reported so far can be traced to avian origins (1918, 1957, 1968, 2009), emergence from wild birds, often facilitated by intermediate hosts such as swine and poultry (Runstadler *et al*., 2013;Worobey *et al*., 2014; Meseko *et al*., 2018). Amplification of IAVs has been well characterized at the domestic animal–human interface. Therefore, these parks and gardens can be a potential source of transmission of HPAI considering the fact that many people visit these gardens and parks and have close contact with these wild aquatic birds. Human infection with certain contemporary Eurasian lineage HPAI H5N1 viruses frequently causes severe disease with a case fatality rate of up to 50%, as reported by Nelli *et al*. (2012). As HPAI continued to spread, the biodiversity of the world’s birds population is threatened as hundreds of thousands of wild birds have died in recent times in an unprecedented outbreak of avian influenza in Europe, North America, and across the globe, raising concerns that these outbreaks will certainly affect the survival of certain species of wild birds. For conservation purposes, it may not be proper to depopulate these types of birds as it is done with domestic poultry. Therefore, it is necessary to put in place effective preventive and control measures through an enlightenment campaign of stakeholders, continuous surveillance and monitoring of these viruses in AIVs at risk areas is advocated.

## Acknowledgments

This study was carried out without any funding. But was supported with all reagents needed in conducting the analysis by the National Veterinary Research Institute (NVRI), Vom, Nigeria. The authors would like to appreciate the staff of the Regional Laboratory for Animal Influenza and Transboundary Animal Disease, NVRI, Vom for the technical support. We also acknowledge Ikemefuna N. for preparing the case report from Delta state.

